# Spatiotemporal reconstruction of emergent flash synchronization in firefly swarms via stereoscopic 360-degree cameras

**DOI:** 10.1101/2020.03.19.999227

**Authors:** Raphaël Sarfati, Julie Hayes, Élie Sarfati, Orit Peleg

## Abstract

During mating season, males of synchronous firefly species flash in unison within swarms of thousands of individuals. These strongly-correlated collective displays have inspired numerous mathematical models to explain how global synchronous patterns emerge from local interactions. Yet, experimental data to validate these models remains sparse. To address this gap, we develop a method for three-dimensional tracking of firefly flashes, using a stereoscopic setup of 360-degree cameras. We apply this method to record flashing displays of the North American synchronous species *Photinus carolinus* in its natural habitat as well as within controlled environments, and obtain the 3D reconstruction of flash occurrences in the swarm. Our results show that even a small number of interacting males synchronize their flashes; however, periodic flash bursts only occur in groups larger than 15 males. Moreover, flash occurrences are correlated over several meters, indicating long-range interactions. While this suggests emergent collective behaviour and cooperation, we identify distinct individual trajectories that hint at additional competitive mechanisms. These reveal possible behavioural differentiation with early flashers being more mobile and flashing longer than late followers. Our experimental technique is inexpensive and easily implemented. It is extensible to tracking light communication in various firefly species and flight trajectories in other insect swarms.

## 1. Introduction

Firefly flashes are more than a mere midsummer night’s wonder: they express a sophisticated social behaviour characterized by male courtship and female mate choice [1]. Firefly swarms are mass-mating events that contain purposeful internal dynamics [2]. Importantly, fireflies offer a rare glimpse into insect communication, as their broadcasting signals, consisting of intermittent and periodic flash patterns, are readily traceable even in congested groups. Therefore, it is possible to separate movement from communication, unlike in other insect swarms where trajectories are a proxy for social interactions [3, 4]. During mating season, male fireflies advertise themselves to stationary females on the ground by flashing their species-specific patterns to be identified as potential mates [2, 5].

In certain species, males flash synchronously in unison within swarms of tens of thousands of individuals [6]. This phenomenon, often reported in Southeast Asia, was first studied by Buck and Buck [7, 8]. In North America, the well-studied colony of synchronous species *Photinus carolinus* in Great Smoky Mountains National Park (GSMNP) mates for 10 to 15 days in early June, a phenomenon that has attracted tourists and scientists alike for many years [9].

The collective flashing displays of *P. carolinus* have been described in detail, notably by Copeland and Moiseff [10, 11], who showed that males flash synchronously every *T*_*f*_ ≃ 0.5s, for bursts of a few seconds, and then collectively stop for a few seconds, leaving their environment completely dark. Flash bursts repeat every *T*_*b*_ ≃ 12 − 14s for up to 3h after sunset and are believed to provide an opportunity for females, located close to the ground, to respond outside of visual clutter [12]. As a landmark of synchronization in nature, these displays have inspired various mathematical models to explain how a number of coupled oscillators might find themselves in sync if given enough time [13, 14]. As illustrated by these models, a comprehensive understanding of firefly collective behaviour requires not only temporal, but also spatial information about flash occurrences, which has been lacking until now. To address this gap, we captured stereoscopic footage of *P. carolinus* flashing displays in GSMNP in order to obtain three-dimensional (3D) reconstructions of flashing swarms.

Traditionally, stereoscopic setups have used regular (planar) cameras for the 3D tracking of flocking or swarming animals, such as mosquitoes [3], midges [4, 15, 16], or birds [17, 18, 19, 20, 21], which have provided considerable insights into the mechanisms involved in collective dynamics, for example the structure of interactions [17, 15, 22] or propagation of information [20, 21]. Over the past few years, however, 360-degree cameras (or 360-cameras) have started to be used widely, notably in computer vision [23]. These cameras stitch together multiple Fields Of View (FOVs) from complementary wide-angle objectives in order to provide a full spherical image. To our knowledge, 360-cameras have not yet been used in the study of swarms and flocks but potentially offer complementing advantages to traditional techniques. Indeed, while the cone-like overlap between two planar cameras’ FOVs is well-suited to small swarms or distant flocks, it suffers from significant limitations for large or extended groups such as firefly swarms, as cameras need to be placed outside the collective dynamics and can only capture a sliver of the action (figure 1*a*). In contrast, 360-degree cameras can be placed directly within a swarm of interest, and present a FOV overlap that is more isotropic (figure 1*a*). For firefly swarms, this even enables the recording of collective male displays from the perspective of a stationary female on the ground.

**Figure 1.**
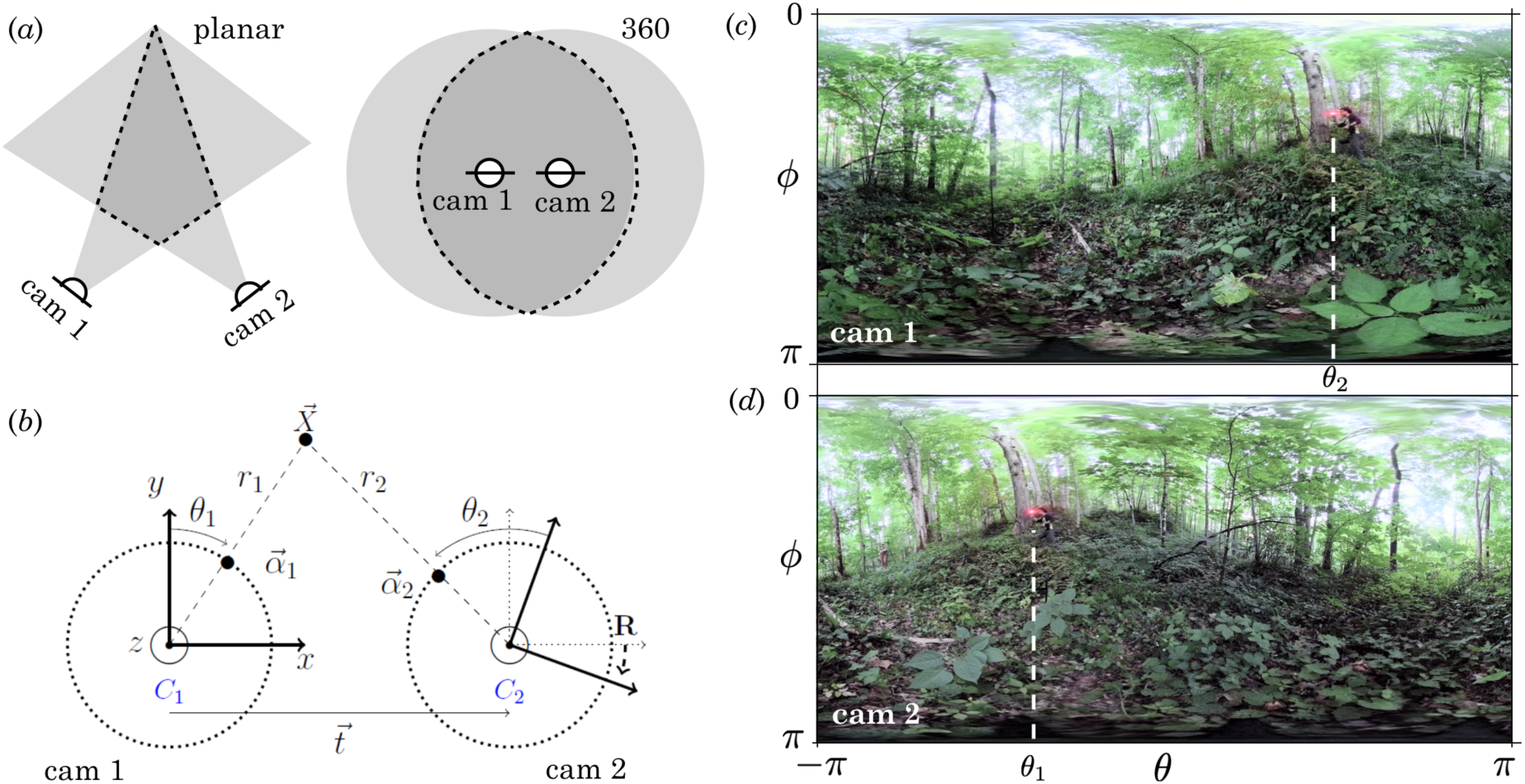
Stereoscopic vision using two 360-cameras. *(a)* Compared to stereoscopic vision using planar cameras, for which the intersection of FOVs is a cone-like shape (left), 360-cameras positioned close to each other have a sphere-like FOV overlap (right). *(b)* Schematic of the principles of 360 stereo reconstruction, showing the position 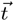 and orientation **R** of *C*_2_ relative to *C*_1_, and the projections of the world point 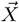 in the FOV of each camera. For simplicity, the schematic assumes only a rotation around the *z*-axis, **R** = **R**_**z**_(*ψ*_*z*_). *(c, d)* To illustrate, FOVs in *(c) C*_1_ and *(d) C*_2_ in equirectangular form. The horizontal coordinate maps onto the polar angle *θ* between *−π* and *π*, and the vertical coordinate maps onto the azimuthal angle *ϕ* between 0 and *π* (top to bottom). The same red dot, representing 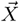, is seen at different (*θ, ϕ*) positions in *C*_1_ and *C*_2_.

Here, we present a general framework for 3D reconstruction using a pair of 360-cameras, and demonstrate its implementation for the analysis of firefly swarms’ internal dynamics, in particular flight patterns and spatiotemporal correlations. This paper consists of three main parts. The next section presents the theory behind the 3D reconstruction technique and its practical implementation, and may be useful to experimentalists for the study of fireflies and other insect swarms. Our Matlab code is provided in the Electronic Supplemental Material (ESM 1.1). The Results section reports findings relative to the behaviour of *P. carolinus* fireflies, and may be of interest to biologists, entomologists, physicists, and even the general public. In the following section, we detail some of the advantages and limitations of our experimental technique, and outline possible applications based on the data presented. We also provide with this paper a standalone FireflyNavigator software tool for the reader to interactively navigate reconstructed firefly swarms and visualize trajectories (ESM 1.2).

## 2. Methods: 3D-reconstruction via pairs of 360-degree cameras

Stereoscopic vision uses image projections from two distinct perspectives to triangulate the positions of world points in 3D. Below, we introduce the model underlying stereoscopic reconstruction from pairs of 360-cameras (epipolar geometry), and detail its practical implementation.

### (a) Theory

#### Single 360-degree camera

A 360-camera can be modeled as a point in space with an internal orientation described by an orthogonal frame 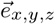 (figure 1*b*). As the camera estimates the angular position (*θ, ϕ*) of a world point 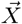 relative to itself, but not its distance, world points are only known with respect to their projections 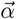 on the unit sphere:

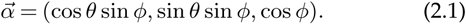

#### Two 360-degree cameras in stereoscopic setup

Consider two 360-cameras, *C*_1_ and *C*_2_. We arbitrarily choose the position and orientation of *C*_1_ as the origin and frame of reference of the world. With respect to *C*_1_, *C*_2_ is translated by a vector 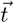, and its internal frame is rotated by three Euler angles (*ψ*_*x*_, *ψ*_*y*_, *ψ*_*z*_), which can be represented by a 3 × 3 rotation matrix **R** = **R**_**x**_(*ψ*_*x*_) · **R**_**y**_(*ψ*_*y*_) · **R**_**z**_(*ψ*_*z*_) (figure 1*b*), where det(**R**) = 1 and **R**^−1^ = **R**^**T**^. Since there is no intrinsic length-scale in this geometry as cameras only evaluate point projections, we set 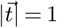, and the correspondence with real-world units can be made by measuring the distance between the two cameras in the experimental setup.

#### Triangulation: theory

Given a point 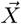 in space, its coordinates are: 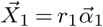 in the frame of *C*_1_, and 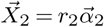 in the frame of *C*_2_. To express 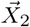 in the frame of *C*_1_, it needs to be rotated back, and therefore has coordinates **R**^−1^ 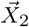. From there, we obtain the geometric relation (vector addition):

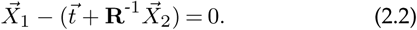

As a consequence, vectors 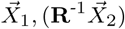, and 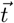 are coplanar, and so are their angular projections, such that

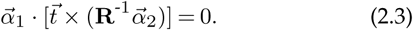

By writing the cross product as a matrix multiplication 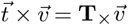, we define the Fundamental Matrix as **F** = **T**_*×*_ · **R**^−1^ [23], such that

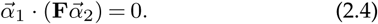

### (b) Implementation

#### Camera data

360-cameras have internal or external software that performs the stitching between the FOVs recorded from different objectives. One common output of these stitching procedures is a movie consisting of equirectangular frames of dimension *n*_*p*_ × 2*n*_*p*_ pixels^2^ which map the planar coordinates (*x, y*) onto the azimuthal angle *θ* and the polar angle *ϕ*: (*x, y*) ∈ ⟦1, 2*n*_*p*_**⟧** × ⟦1, *n*_*p*_**⟧** −→ (*θ, ϕ*) ∈ [0, 2*π*[×[0, *π*] (figure 1*c,d*). The angles are obtained using:

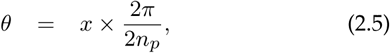

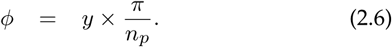

and the spherical coordinates 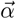 are defined in Eq. 2.1. The 3D reconstruction from sets of points {(*θ, ϕ*)} in pairs of frames requires three steps: calibration, matching, and triangulation, as described in the following paragraphs.

#### Calibration

Calibration aims to determine the position 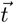 and orientation **R** of *C*_2_ relative to *C*_1_. We assume that we have a set of *N matched* points between the two cameras. (These can be obtained from an identifiable trajectory, or using specific points such as the corners of a checkerboard, or by manual identification of specific features.) From the set of matched points, we propose two methods to compute 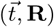.

Method 1: optimization search. The idea is to find the set of 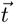 and **R**-coordinates that minimizes the sum of the distances between triangulated points in each camera. Assuming test values 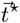 and **R**^*\**^, there exists a set of distances *r*_1_, *r*_2_ ≥ 0 such that 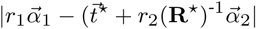 is minimum [23]. The optimization search occurs on a 6-dimensional space since 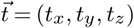 and **R** is determined by the three Euler angles ψ_*x,y,z*_, with the additional constraint 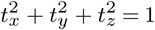. Finally,the problem reads:

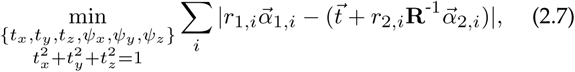

and can be solved using Matlab’s `fmincon` function.

Method 2: fundamental matrix. The camera pose can also be determined from the fundamental matrix, which can be computed using the Matlab built-in function `estimateFundamentalMatrix`. (Note that this function is designed for planar calibration, and therefore it only takes 2D vectors as arguments, assuming that the third coordinate is 1. Consequently, using this function requires renormalizing the projection vectors 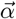 by their third coordinate 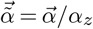.) From the estimation of **F** = **T**_*×*_·**R**^−1^, 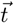 is simply a unitary vector in the null space of **F**^−1^, which leaves an ambiguity of factor ±1. Since det(**T**_*×*_) = 0, estimating **R** relies on singular-value decomposition, for example using the method described in Ref. [24]. Each possibility for 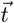 contains two possibilities for **R**, which makes a total of four solutions. Of those, two are improper rotations (determinant -1), which leaves only two possibilities. Choosing the right one may rely on an experimental estimation; for example, it is convenient to arrange the cameras so that **R** ≃ **I** (identity matrix).

Both methods are susceptible to numerical imprecision. For Method 1, optimization search may return a local (rather than global) minimum. For Method 2, the fundamental matrix factorization has more than one solution, and singular-value decomposition can have large numerical errors for ill-conditioned matrices. Therefore, we recommend that both methods be employed to verify that they provide consistent results (ESM 6.2). If they do not, where and why they fail should be examined.

#### Matching

Points extracted from pairs of frames are originally not matched, so a pairing algorithm has to be employed to determine point correspondences. Given 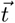, **R**, a set of *n*_1_ points 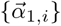 in *C*_1_, and *n*_2_ points 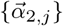 in *C*_2_, an optimal pairing can be made by applying the Hungarian algorithm on an appropriate cost matrix [*c*_*i,j*_]. We chose the cost of pairing 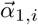 with 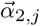 as equal to the smallest possible distance: 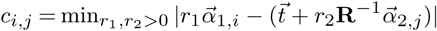.

#### Triangulation: implementation

Given the camera pose 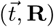 and two matched points 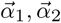 corresponding to projections of world point 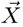, the distances *r*_1_, *r*_2_ ≥ 0 from *C*_1_, *C*_2_ respectively can be calculated by minimizing the distance 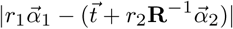 using a linear solver such as Matlab’s lsqnonneg. From there, 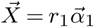. Note that triangulation is impossible for points lying on the same line as 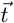. Individual 3D points can be subsequently concatenated into streaks (spatial localization of a single, continuous flash) and trajectories (sets of consecutive streaks from the same individual), as described in the ESM (Section 5).

## 3. Experimental Setup

Data collection on the firefly species *Photinus carolinus* took place in early June 2019 on the Tennessee side of GSMNP per research permit (GRSM-2019-SCI-2075) by the National Park Service. We performed two types of experiments: recordings in the fireflies’ natural habitat, and recordings in a controlled environment.

### (a) Equipment

We used two GoPro Fusion cameras as our 360-cameras. Temperature was recorded using two Kestrel Temperature Data Loggers (one reading every 5min). The cameras were positioned on small tripods (0.6m above ground), and aligned manually as precisely as possible to have the same side facing the same direction (so that **R** is close to identity). The spacing between the two cameras was always set to 3ft (0.91 m) using a wooden yard stick (figure 2*b*). We recorded either at 30 or 60 frames-per-second (fps), and the ISO was manually set to 1600. We applied black electrical tape on the screens and LEDs of the cameras so as not to perturb fireflies with artificial light signals. The recorded footage was then processed using the software provided with the cameras, GoPro Fusion Studio, in order to create equirectangular movies in high-resolution (4K) MPEG format which could later be processed in Matlab (figure 1*c,d*). It is crucial to render the movies using no stabilization option in order to maintain constant orientation throughout the movie. In order to identify simultaneous frames in both cameras, a brief light signal was triggered a few seconds after recording started. The beginning of the signal marked the frame of reference in each movie, allowing us to estimate the exact delay (within one frame) between cameras. We later found that using cross-correlations between frames from both cameras resulted in identical delay estimations, and used that to confirm that delays remain constant even after 2 hours of recording. Calibration was performed using the trajectory of a small LED. For camera pose estimation, we used the results from the fundamental matrix computation after verifying consistency with the other proposed methods (see ESM 6.2).

**Figure 2.**
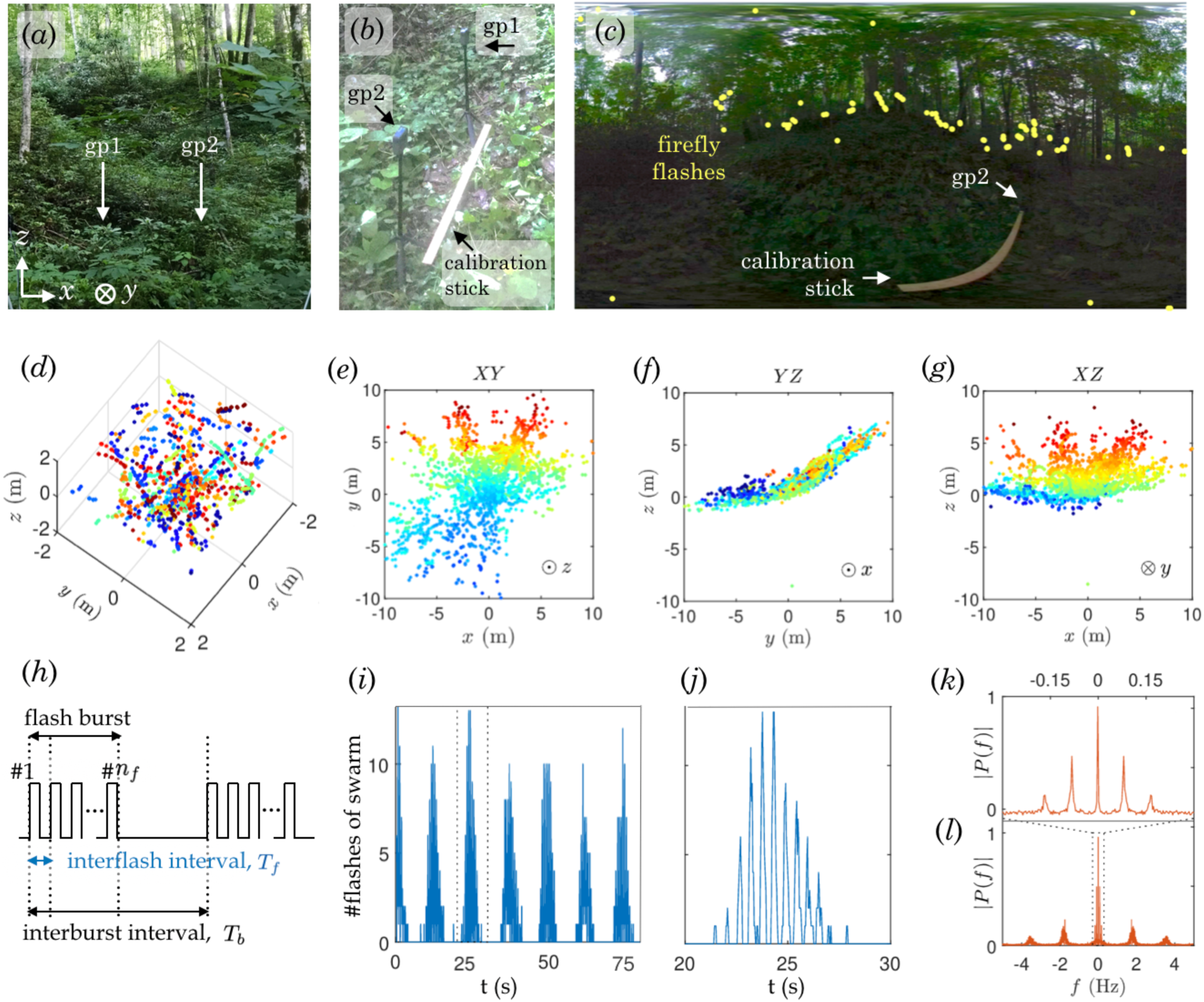
Spatial and temporal patterns in field experiments. *(a)* Broader view of GoPro surrounding environment. The rising ridge is clearly visible in the distance. The cameras (gp1 and gp2) are situated approximately as indicated, but surrounding vegetation conceals them. The coordinate system is defined in the bottom-left corner, with *xy* defining the horizontal plane, and *z* the vertical axis. *(b)* GoPro cameras standing on small tripods and separated by 0.9m. *(c)* Contrast-adjusted 360-view from gp1, with yellow dots showing the locations of a few firefly flashes, mostly concentrated along the ground. The yard-stick in *(b)* is also seen, with gp2 standing at the other end. See also Movie S1. *(d)* 3D reconstruction in a 2 *×* 2 *×* 2m^3^ cube, centered around gp1. Colors indicate occurrence in time (blue to red), over 5min. See also Movie S2. *(e, f, g)* 2D projections of the full reconstructed swarm, from *(e)* above, *(f)* the side, and *(g)* the front. Colors (blue to red) indicate value along the axis perpendicular to the page (*z, x, y*, respectively, as indicated in the bottom right corner of the plots). *(h)* Schematic of *P. carolinus*’s flash pattern. Flashes are produced in bursts of variable *n*_*f*_ flashes. These bursts are separated by a second time-scale, the interburst interval *T*_*b*_ (time from the onset of one flash burst to the onset of a consecutive flash burst). During the time between bursts, no flashes are produced. *(i)* Time series of number of flashes per frame over 2min30s. Bursts of collective flashing occur at regular intervals (about 12.5s). *(j)* Zoom on the flash burst between *t* = 20s and *t* = 30s. The burst shows a succession of peaks at regular intervals (about 0.5s), suggesting synchrony. The characteristic triangular shape of the burst is seen for all bursts. *(k,l)* Fourier transform of the time series in *(i)* showing two distinct frequencies. The low frequency in *(j)*, about 0.08Hz, corresponds to interburst intervals. The high frequency in *(l)*, about 1.75Hz, corresponds to interflash intervals.

**Figure 3.**
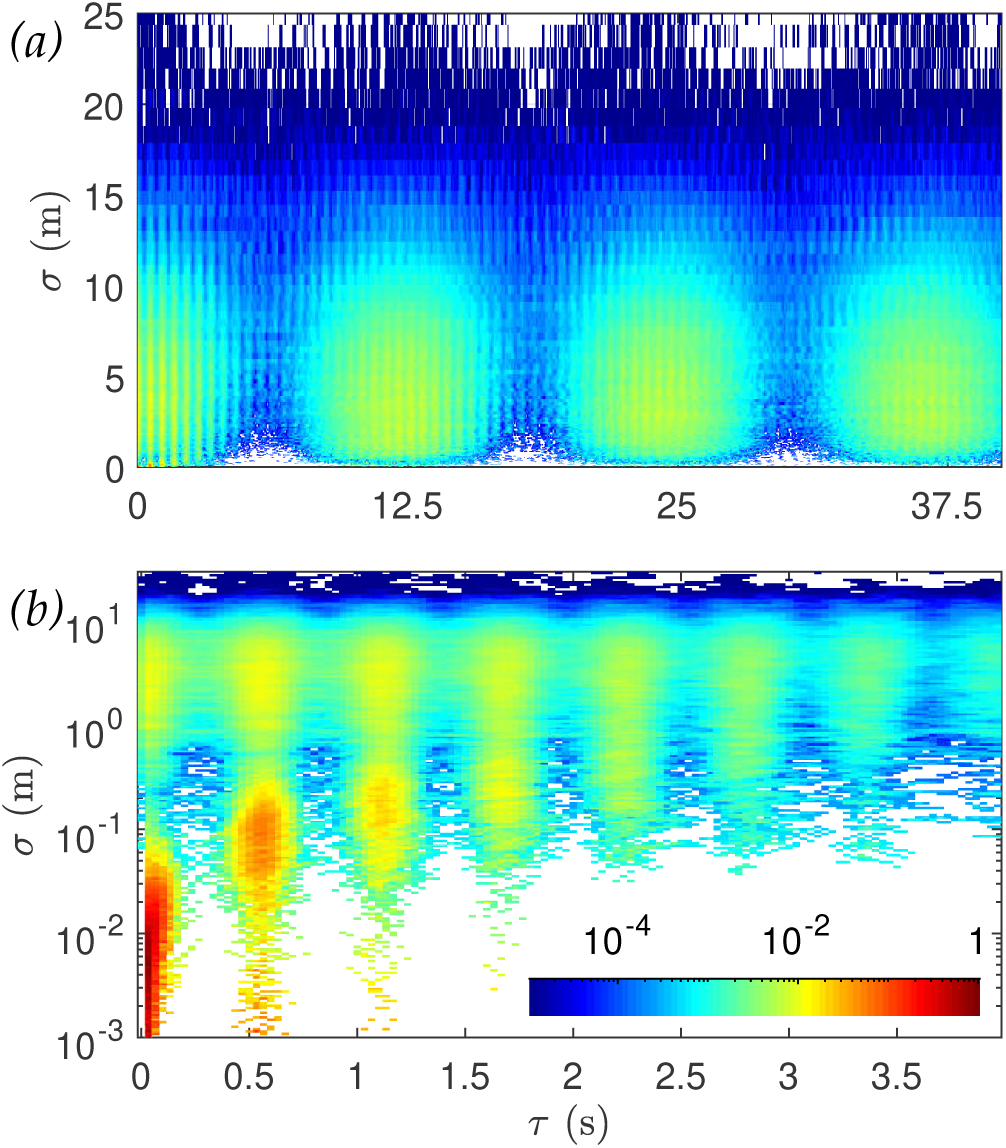
Spatiotemporal correlations: probability distribution functions in the delay-separation (*τ, σ*) plane. *(a)* Large time- and length-scales, showing interburst correlations. *(b)* Small time- and length-scales, showing intraburst correlations.

**Figure 4.**
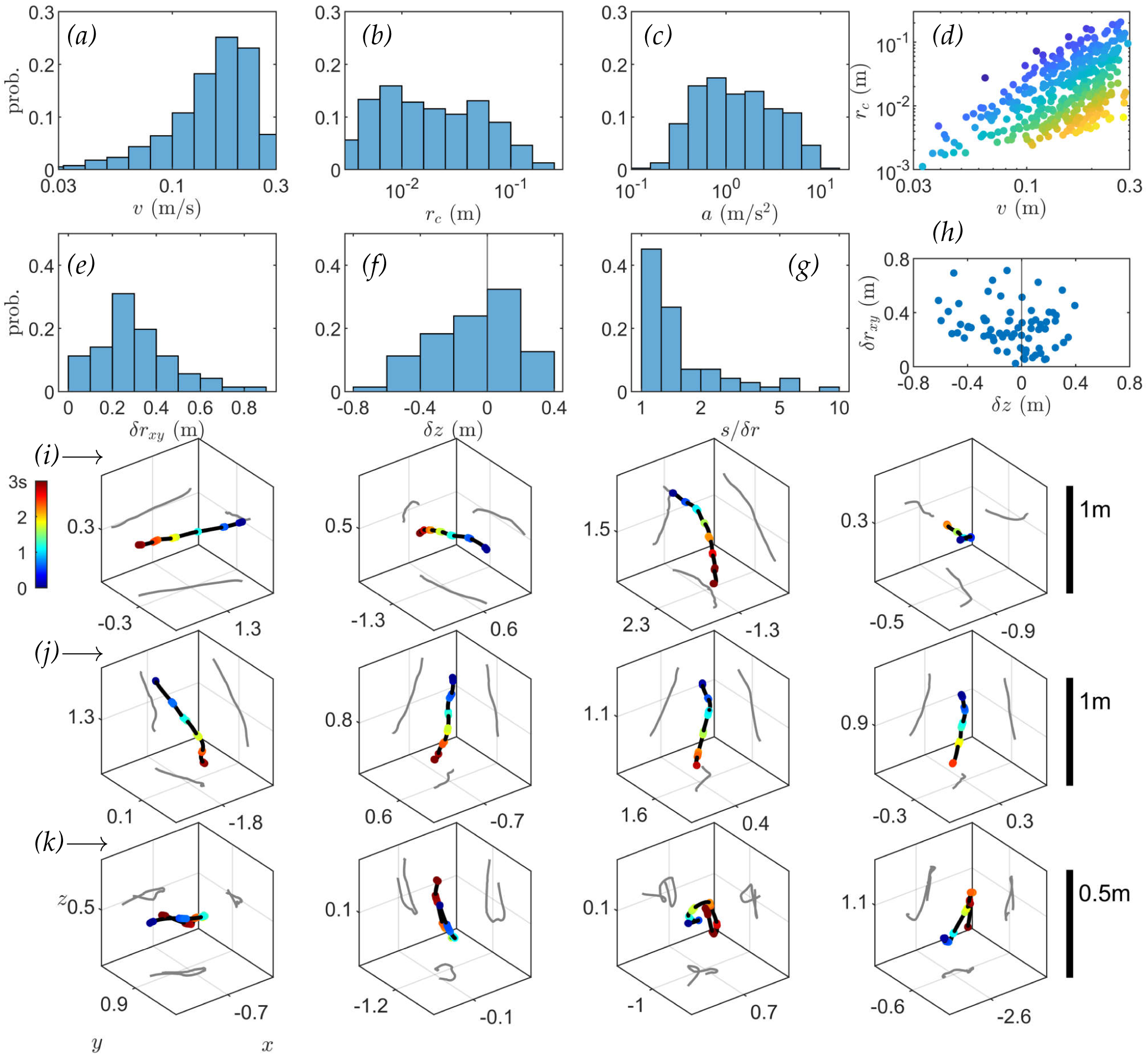
Kinematics of natural flights. *(a-c)* Distributions of *(a)* streak velocity *v* (mean frame-to-frame displacement), *(b)* radius of curvature *r*_*c*_ (circle fit to streak positions), and *(c)* acceleration *a* = *v*^2^*/r*_*c*_ for naturally-occurring firefly flashes. *(d)* Scatter plot of *r*_*c*_ vs *v*, with colours indicating *a* value (blue to yellow). Accessible kinematics span only a portion of the plane; notably, the high-acceleration regime (bottom-right; fast and curved streaks), and the slow-straight regime (top-left) are excluded. *(e-g)* Trajectory (ensemble of streaks belonging to the same firefly) metrics: distributions of *(e)* horizontal displacements *δr*_*xy*_, *(f)* vertical displacements *δz*, and *(g)* trajectory length *s* over end-to-end distance *δr. (h)* Scatter plot of vertical displacements vs horizontal displacements. *(i-k)* Representative trajectories for three different regimes: row *(i)* large horizontal displacements, row *(j)* large vertical displacements, row *(k)* long trajectories with close end-points. Coloured points indicate recorded (triangulated) flashes, and black lines interpolated paths. Rows *(i)* and *(j)* show cubes of 1 *×* 1 *×* 1m^3^, and row *(k)* 0.5 *×* 0.5 *×* 0.5m^3^, for scale, with axes value indicating mean position with respect to gp1.

### (b) Data collection in natural habitat

The specific GSMNP site for natural habitat recordings was situated between a trail along a creek and a steep ridge, in a bushy area (figure 2*a*) that had been observed in previous nights to show high activity of *P. carolinus*. Prior to the start of the display (about 30min before sunset), the two 360-cameras were placed in a small terrain depression clear from trees, and close to the bottom of the ridge (figure 2*b*). They were positioned side by side on firm ground, and facing the same direction (figure 2*b*). Recording at 30fps was started using a remote control at 9:15pm EST, 29min after sunset (8:46pm), and continued for about 90min. Local ambient temperature was 18.5 ± 0.5 °C.

### (c) Data collection in controlled environment

In addition to recording firefly displays in the natural, unperturbed habitat, we performed a series of controlled experiments, in which a specific number of *P. carolinus male* fireflies were placed in a large tent. For these experiments, fireflies were gently captured during peak flashing hour using insect nets, and delicately placed into petri dishes for up to a few minutes before being introduced into the tent, where they were visually inspected to confirm their sex and species (see ESM 3.1). They were then placed by increasing numbers into a cuboid black fabric tent of dimensions 2m×1.5m×1.5m (*x*-*y*-*z*, figure 5*b*). An additional black plastic tarp was added on top in order to ensure visual insulation from fireflies on the outside. Outside temperature decreased from 19*°*C to 17*°*C over the course of the experiments, while the temperature inside the tent was 18*°*C. Each experiment lasted for 15min recorded at 60fps (about 55000 frames), and consisted of the same stereoscopic setup (figure 5*a*).

**Figure 5.**
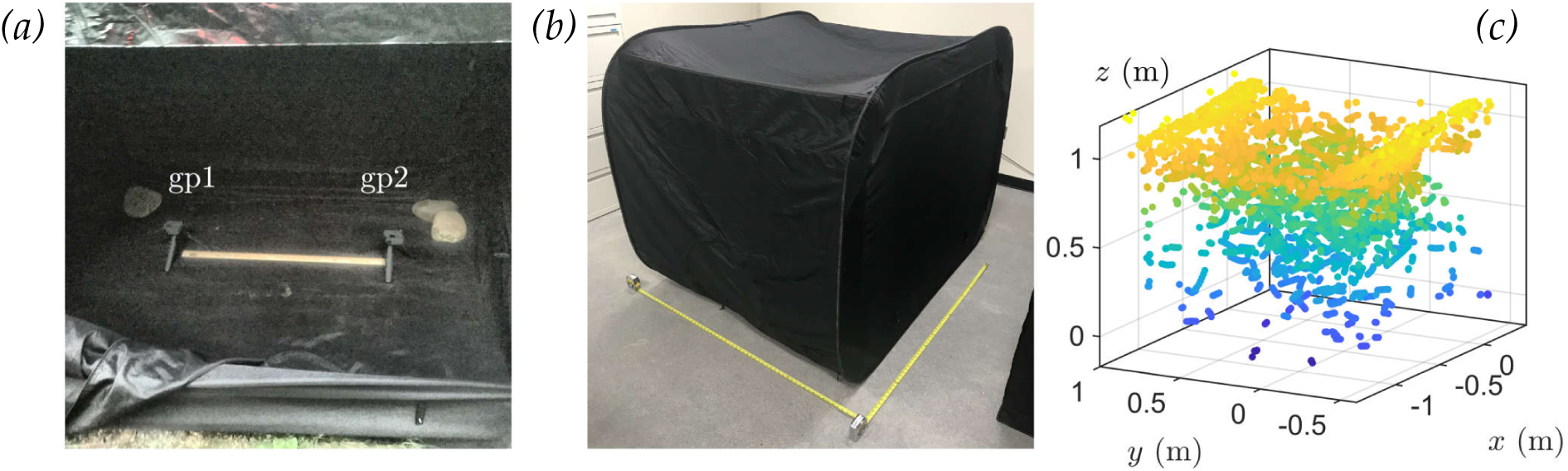
Controlled experiments in a confining tent. *(a)* GoPro cameras inside the tent, separated by the yard stick (0.9m). *(b)* Tent geometry. The yellow tape measure indicates 1.5m. The *x*-side on the right is about 2m long, and contains the zipper opening seen in *(a)*. The *y*-side and *z*-side (vertical) are about 1.5m-long. Note the roof fabric curves under its own weight. *(c)* 3D reconstruction of flash occurrences in the tent (colors indicate height, from blue to yellow). The dimensions of the enclosing volume match tightly those of the tent, and the shape of the top layer mirrors the curvature of the tent’s roof.

In the set of experiments presented here, we introduced a single firefly first, and additional fireflies subsequently every 15min to reach cumulative numbers of *n* =5, 15, 25, and 40. Due to time constraints and the difficulty of finding fireflies in the tent, we did not attempt to remove fireflies before introducing new ones. All fireflies were released after no more than two hours, and great care was taken so as not to harm them.

## 4. Results

### (a) Natural habitat

#### Spatial distribution of flash occurrences

The 3D reconstruction of flash occurrences in its natural habitat (5min-interval starting at 10pm) shows a *P. carolinus* swarm that closely follows the slope of the surrounding terrain, and notably flashes almost exclusively in a layer of about 2m above ground (figure 2*d-g* and Movie S2). Viewed from above, the reconstructed swarm reveals the limits of the imaging technique: flashes farther than about 10m are not captured, and visual occlusion creates significant “blind zones”. However, triangulated positions show clear streaks of lights (figure 2*d*). This dataset is available for visualization with FireflyNavigator (ESM 1.2).

#### Temporal pattern of flash occurrences

In every frame of the movie, zero, one, or several flashes are captured. The time series of the number of flashes is presented in figure 2*i,j* and shows a doubly periodic pattern. Bursts of flashes happen at regular intervals (*interburst* intervals *T*_*b*_, figure 2*i*), with a maximum of about 15 simultaneous flashes recorded, and are separated by periods of absolute darkness. By zooming on these bursts (figure 2*j*), another temporal pattern appears: bursts consist of a train of a few flashes, happening synchronously, also at a well-defined *interflash* interval *T*_*f*_ *≃* 0.5s. The frequency spectrum (Fourier transform) of the flash time series further confirms the regularity of these two processes by revealing pronounced peaks at frequencies 1*/T*_*b*_ =0.08 Hz and 1*/T*_*f*_ =1.75 Hz (periods of 12.5s and 0.57s, respectively; figure 2*k,l*). The fact that these frequencies appear as sharp peaks in the power spectrum indicates that these two processes occur at well-defined time intervals. These simple quantitative results demonstrate that the *P. carolinus* flashing display is synchronous, intermittent, and precise, in agreement with previous measurements by Copeland and Moiseff [10, 11] which describe similar intermittent (or “discontinuous”) synchrony. However, unlike their previous observations that “group flashing terminated abruptly” [10], we consistently observed a triangular shape of flash bursts, with a slow fading-out phase over a few beats (figure 2*j*). This triangular pattern might suggest some underlying propagation of information within the swarm (ESM 4), and is therefore an important feature of collective flashing. In controlled experiments described below, we show that this shape is not an experimental artifact.

#### Spatiotemporal correlations of flash occurrences

These results demonstrate that a swarm of *P. carolinus* males is a strongly correlated system. The mechanisms underlying their collective behaviour, such as information propagation, can be uncovered by the study of spatio-temporal correlations. For each recorded flash occurrence, we associate a time *t*_*i*_ and a 3D position 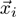. Then, for every pair of flash occurrences (*i, j*), we calculate the separation 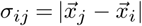 and the delay *τ*_*ij*_ = |*t*_*j*_ − *t*_*i*_|, and we consider the distribution of *σ* vs *τ* in figure 3. The probability density is displayed by colour using a logarithmic scale, and the smallest increment for *τ* is 1 frame (0.033s). The temporal structure of flash occurrences, as reported in figure 2, is reflected in this distribution: correlated peaks occur every 12.5s (figure 3*a*), corresponding to flash bursts, and each of them consists of a series of high and low **5** fringes every 0.55s (figure 3*b*), corresponding to interflash intervals. Spatial correlations between bursts in figure 3*a* extend across the entire swarm (peak in the 0-10m range), demonstrating that flash bursts span the entire (recorded) swarm. Spatial correlations at short times (figure 3*b*) exhibit a bimodal distribution along the *σ*-axis. The peaks at small *σ* correspond predominantly to correlations within a streak (*τ* ≤ 0.1s) and between successive streaks from the same firefly (*τ* = 0.55, 1.10, 1.75, … s). The peaks in the 1-10m range, at all delays including *τ* = 0s are more significant, and suggest that there is no characteristic timescale for information propagation, at least at resolvable times. This important result, which will be investigated in more depth in future work, could be well-explained by the following hypothesis: due to significant visual occlusion, and a mixture of fireflies at rest and moving, it is possible that information transfer relies on a network of visual connectivities with no well-defined length-scale. Two fireflies at short distance might not be able to interact due to visual occlusions, but two fireflies far apart could if connected by a line of sight.

#### Flight kinematics

3D reconstruction also provides insights into the kinematics of moving fireflies. The analysis of individual streaks (flashes spanning at least 4 consecutive frames) shows a wide range of fireflies’ motilities. Streak velocities *v* show a continuum between immobility and fast flights at speed up to 30cm/s (figure 4*a*; the distribution reported here is expected to contain artifacts as flying flashers are more likely to be recorded than immobile ones, which suffer from greater visual occlusion). Comparable, but usually larger, velocities have been observed in other insect flights [25, 26]. Streak curvature radii *r*_*c*_ are also widespread, revealing sharp turns as well as straight flights (figure 4*b*). Streak accelerations, calculated as *a* = *v*^2^*/r*_*c*_, span two orders of magnitude, with an upper limit comparable to the Earth’s gravity (figure 4*c*), analogous to what has been seen in other insects [25]. Interestingly, the distribution of *r*_*c*_ vs *v* shows two well-defined limiting branches (figure 4*d*). The lower branch (large *v*, small *r*_*c*_) marks the “high-acceleration” regime, corresponding to sharp and fast turns. The upper branch is more surprising, and suggests that slow and straight trajectories are not possible per firefly propulsion.

Next, we report observations pertaining to long recorded trajectories (longer than 2s), typically consisting of at least 4 streaks. These trajectories show different patterns. Considering the horizontal excursion *δr*_*xy*_ between trajectories’ end-points, we observe in figure 4*e* a continuum between trajectories which are almost stationary, and others that cover up to 1m. The vertical excursion *δz* is asymmetrically distributed (figure 4*f*), with downward trajectories typically extending farther. Trajectories appear to be never completely vertical (large |*δz*|, small *δr*_*xy*_) but sometimes completely horizontal (figure 4*h*), which potentially suggests limitations of flight capabilities. Furthermore, the ratio of a trajectory’s path length *s* to its end-to-end distance 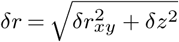 (figure 4*g*) shows that while most trajectories are rather straight (*s/δr ≃* 1), a significant fraction seem very curved and loopy (large *s/δr*).

For illustration, we present a few trajectories corresponding to these different cases: large horizontal excursions (figure 4, row *h*), large downward displacements (row *i*), and highly curved (row *j*). These trajectories are shown in boxes of (1m)^3^ (rows *h, i*), or (0.5m)^3^ (row *j*) for scale. These different types of trajectories may be hypothesized to correspond to different stereotyped behaviours. For example, long and downward trajectories have been observed in male fireflies courting a responding female near the ground [27]. Large horizontal excursions might correspond to exploratory phases.

### (b) Controlled environment

A known number of *P. carolinus* males were placed in a tent in order to study flashing interactions among a small number of fireflies.

#### 3D reconstruction

The 3D reconstruction of flash occurrences in the tent over 15min at *n* = 40 is shown in figure 5*c*. Aside from a small fraction of points (about 1%) which were localized far above the others and were removed from the figure (see discussion in the ESM, Section 6.2), the triangulated points define a volume which closely resembles the tent’s geometry. In particular, dimensions are consistent, and the outline of the curved roof appears clearly (the roof’s fabric curves under its weight, figure 5*b*), with a concentration of points at the edges. In accordance with visual observations when emptying the tent at the end of experiments, fireflies tend to stand on the roof and walls, and hide in the edges and corners, especially in the sharp angles at the junction between the roof and the walls.

Although the confining conditions of the tent plausibly perturbed natural behaviour to some extent, the accessible volume was large enough (∼ 4m^3^) that fireflies were able to move and fly freely, as seen in the trajectories in figure 6*c1-4*. This dataset is available for visualization with FireflyNavigator (ESM 1.2).

**Figure 6.**
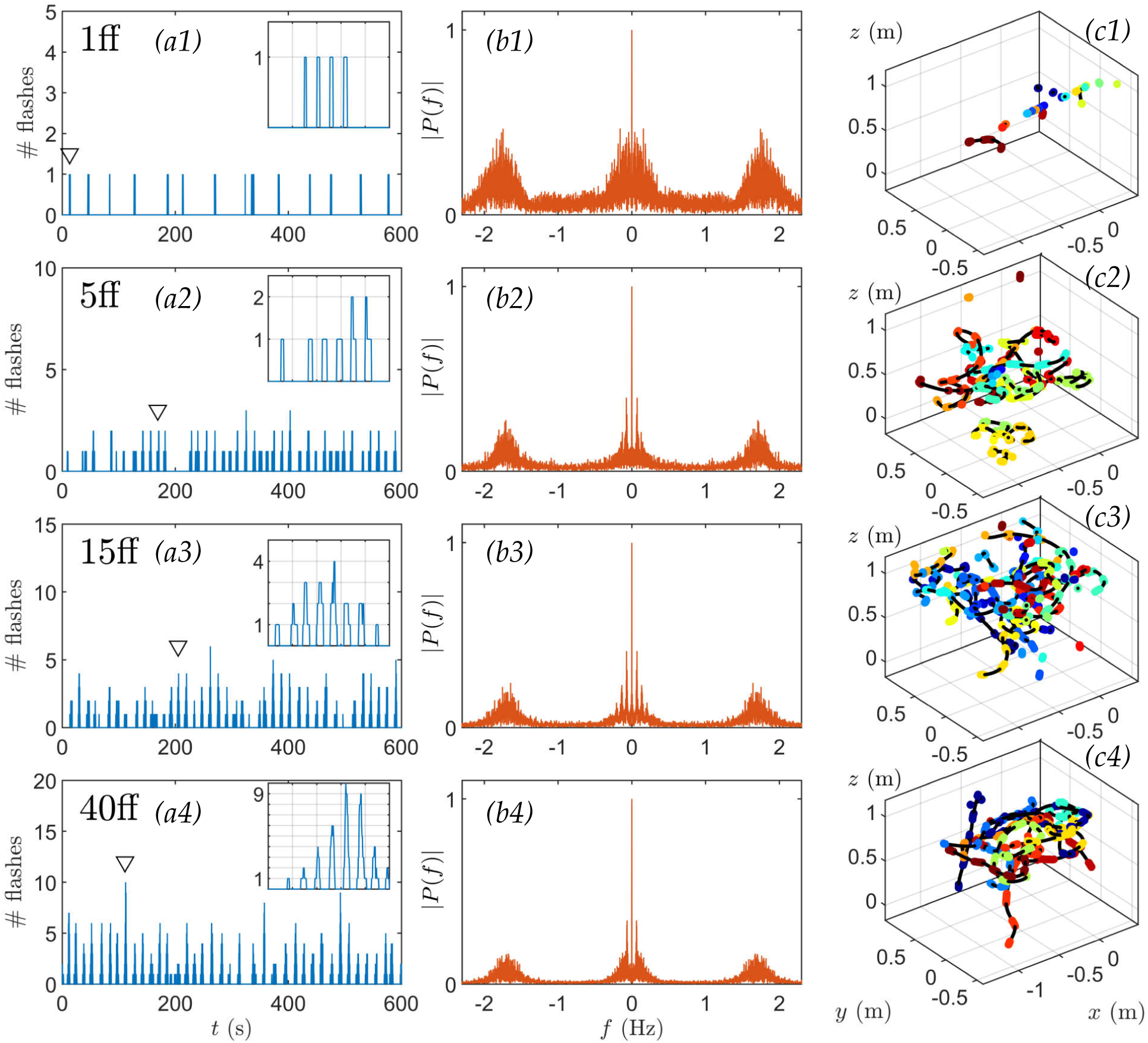
Emergence of collective behaviour in controlled experiments. *(a1-a4)* Time series of the number of flash occurrences over 10min recorded in the tent containing 1,5,15, and 40 fireflies, respectively. Insets: close-up of single bursts (over 5s) indicated by downward triangles. *(b1-b4)* Corresponding frequency power spectra |*P* (*f*)| (Fourier transforms), showing the interflash frequency at 1.75Hz and the emergence of a burst frequency at 0.08Hz when several fireflies interact. *(c1-c4)* Example trajectories (longest flights) for corresponding confined fireflies. Colours indicate time over 15min (blue to red).

#### 1 firefly

When a single male firefly was introduced in the tent, it emitted flashes continuously over the 15min of the experiment (figure 6*a1*), even in the absence of a responding female. Flashes lasted typically between 0.10s and 0.15s (5 to 10 frames), although shorter and longer flashes were also recorded (figure 7*a*). Flashes occurred by bursts of typically 4 consecutive flashes, and overall between 1 and 6 (figure 7*b*), and independently of the pattern, the time interval between two successive flashes was sharply distributed around 0.45s (25 to 30 frames), as evidenced in both the distribution of interflash intervals (figure 7*c*) and the 1.75Hz-peak in the frequency spectrum (figure 6*b1*). Flashing occurred while both flying or standing on the tent’s structure (figure 6*c1*). These observations are generally consistent with previous studies of *P. carolinus* flash patterns [10, 11]. Most importantly, unlike interflash intervals, time intervals between successive bursts did not show any regularity, spanning a wide range from 12s to 1min (figure 7*d*). Similar findings occurred in repeated experiments (see ESM 3.2).

**Figure 7.**
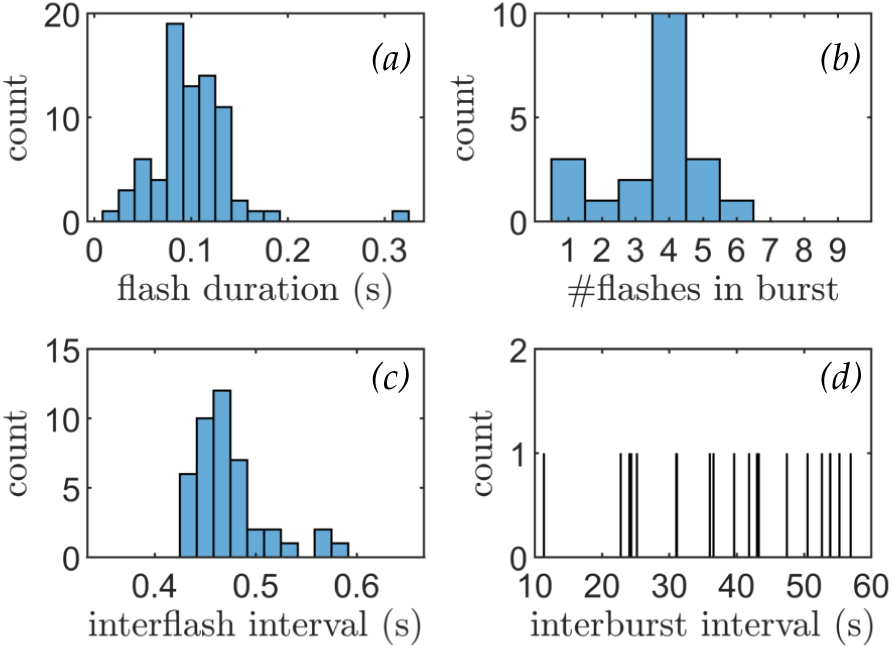
Statistics of a single firefly’s flash pattern, recorded at 60fps. *(a)* Distribution of flash durations. Most flashes last between 5 and 10 frames (0.08s to 0.16s). *(b)* Distribution of number of flashes per burst. A typical burst consists of 4 flashes, although shorter or longer ones are also common. *(c)* Distribution of interflash intervals. Flash intervals appear as very regular, at 0.45*±*0.05s. *(d)* Distribution of time intervals between successive bursts. There seems to be no characteristic time between two bursts, unlike what is observed in collective flashing.

#### 5 fireflies

Four fireflies were subsequently introduced to bring the total to *n* = 5. Flashing continued throughout the experiment, with many flights recorded (figure 6*a2,c2*). It appears that fireflies attempted to synchronize their flash signals, as evidenced by the temporal distribution of flash occurrences. Indeed, the majority of flash bursts comprised at least two simultaneously active fireflies, whose flashes occurred synchronously (figure 6*a2*). Trajectory identification, enabled by the spatial localization of flash streaks, provides further insights into the onset of collective synchrony. Figure 8 shows that the firefly who initiates a burst tends to flash the longest, and that followers start their own flashes *already synchronized*. This strongly contrasts with common mathematical models which describe the onset of synchrony in coupled oscillators through a distribution of phases which becomes continuously sharper over time [13, 14]. Followers can either stop before the flashing leader (figure 8*b*), or continue after him (figure 8*a*), which suggests that flashing information could be transferred in a relay-like manner throughout large swarms. Finally, while flashing bursts seem aperiodic (*e*.*g*., large gap at *t* = 200s in figure 6*a2*), the emergence of a peak at low frequencies (figure 6*b2*) hints at some regularity in the collective flashing pattern.

**Figure 8.**
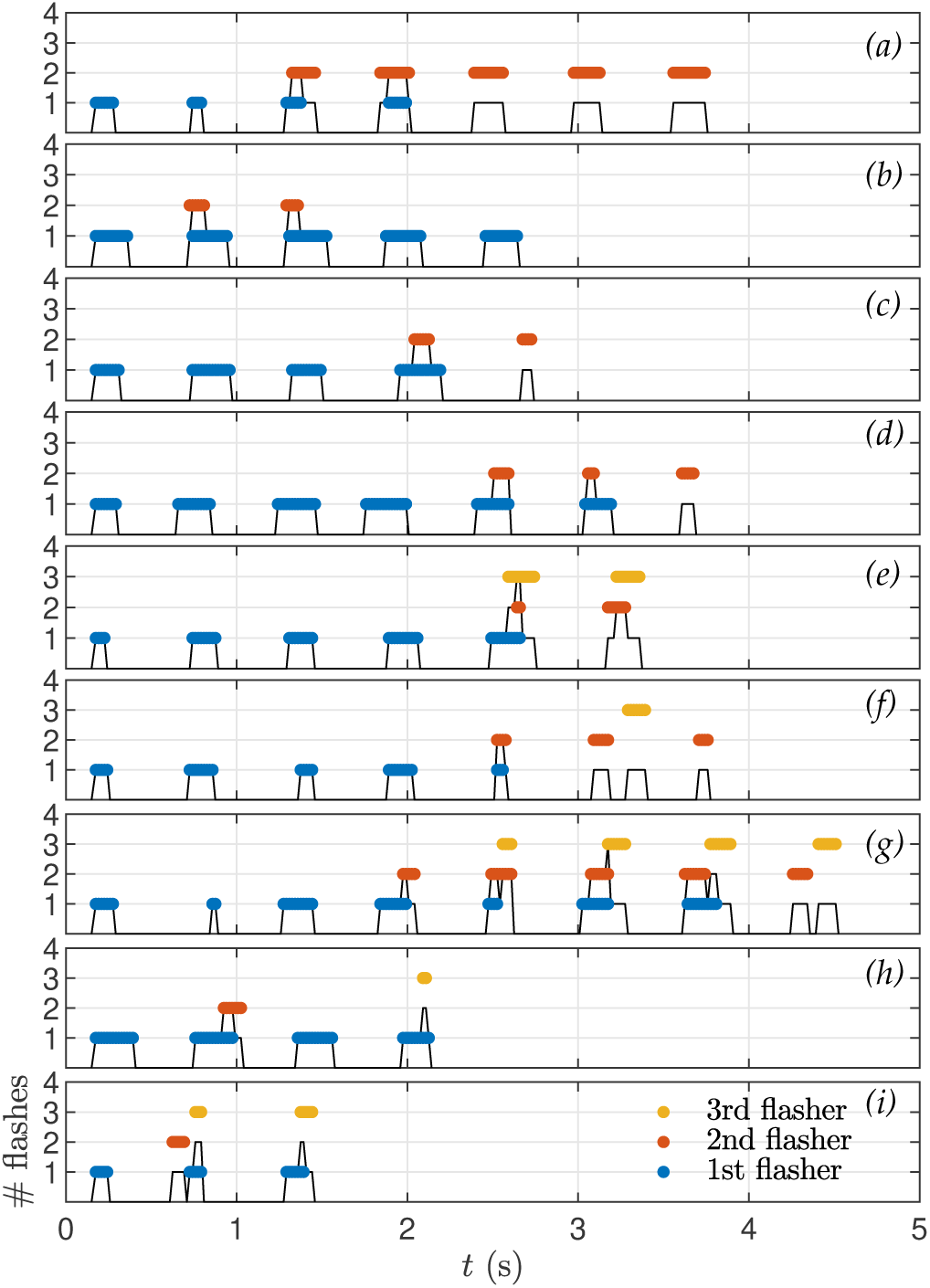
Patterns of synchronous collective flashing. The black line indicates the number of flashes at a given time. The colored segments indicate flashes coming from the same firefly. In panel *(a)*, for example, one firefly starts flashing, and repeats 3 more times (blue). A second firefly starts on the 3^rd^ beat, and continues for a total of 5 flashes. It is slightly delayed on the 3^rd^ beat, but starts first on the 4^th^. In panel *(e)*, a second and third firefly start together (although slightly late) on the 5^th^ flash.

#### 15 fireflies

This regularity becomes more pronounced at *n* = 15, where bursts occur periodically in the time series (figure 6*a3*) and prominent peaks (and their harmonics) emerge in the frequency spectrum at 1*/T*_*b*_ = 0.08Hz (figure 6*b3*). This interburst frequency is identical to the one measured in the wild, and was absent in the flash pattern of a single firefly. Therefore, this suggests that occurrence of a well-defined interburst interval is an emergent property of collective behaviour. The interflash interval at 1.75Hz remains similar to a single firefly’s (figure 6*b3*).

A second important observation concerns the collective kinematics during a burst. In most bursts, only one firefly is seen flying, while others are standing or walking (figure 9, ESM 3.3, and Movie S3). The flying trajectory typically starts the earliest, and comprises the most flashes. This observation could be related to a mechanism which optimizes information transfer while conserving the group’s collective energy resources. Alternatively, it could reveal behavioural differentiation. It was not possible to determine whether it is always the same firefly that is flying in different bursts, as too much localization information is lost in the few seconds of darkness between bursts.

**Figure 9.**
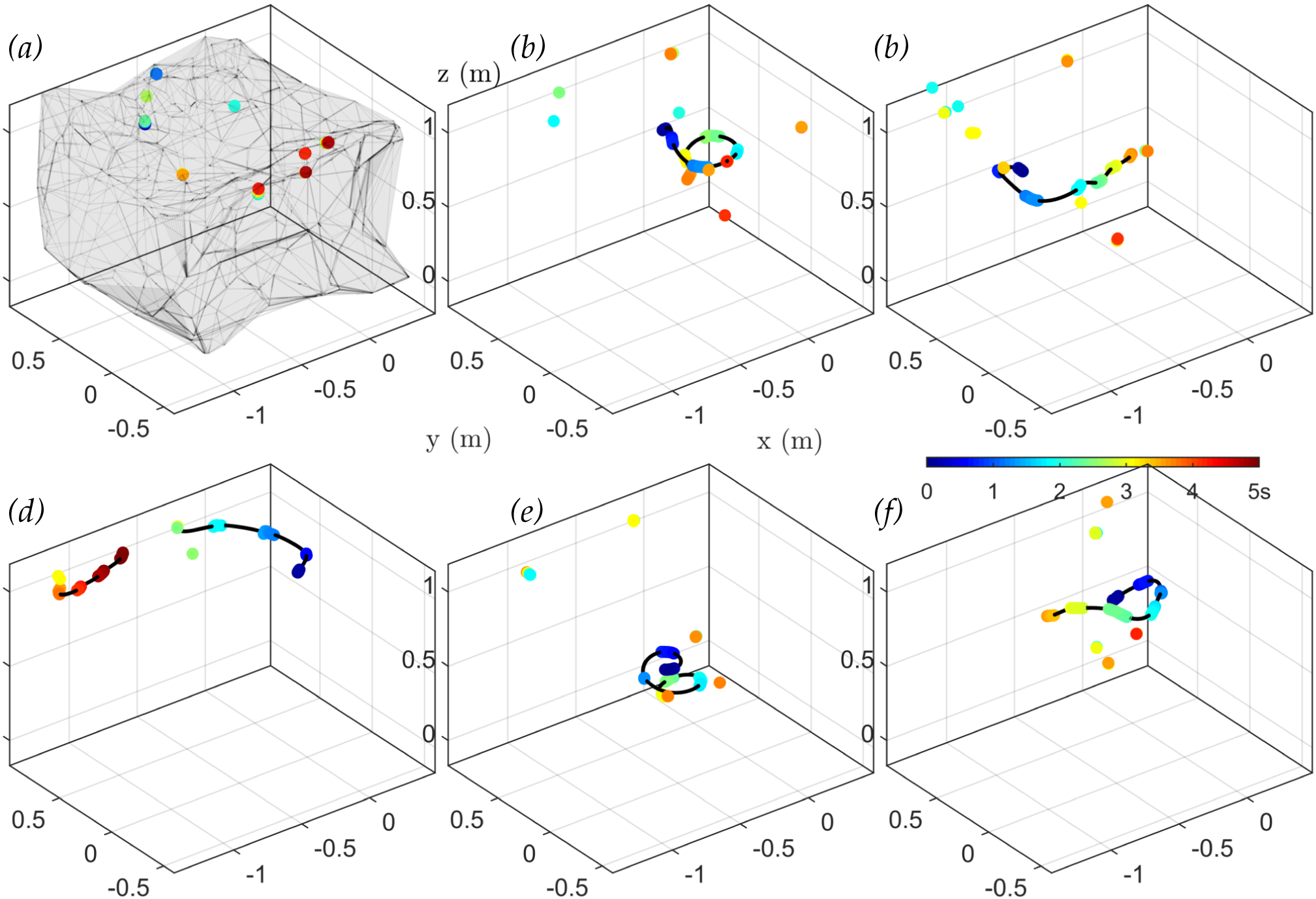
Firefly trajectories during flash bursts. Coloured dots indicate recorded flash occurrences over a time window of 5s (blue to red), and black lines indicate interpolated flight paths. The gray volume in the first plot represents the approximate tent outline obtained from all flash positions (alpha shape), for reference. All 6 plots have the same scale and position. While many flashes occur during a burst, most of them come from immobile fireflies, usually standing on the tent’s ceiling or walls. See also ESM 3.3 and Movie S3.

#### 40 fireflies

The flashing dynamics at 40 fireflies is qualitatively similar to the 15-firefly case, but the larger number of flashes provides more accurate observations. Flashes occur during bursts regularly spread in time; each burst consists of a few synchronous flashes, and presents the same triangular shape as observed in the wild, wherein the number of flashes slowly increases, reaches a maximum, and then slowly decreases (figure 6*a4*). This pattern can easily be considered as the extension of the pair-synchronization presented with 5 fireflies to the case of many fireflies (ESM 4).

In summary, these controlled experiments at increasing density of fireflies show that the synchronous, intermittent flashing display of *P. carolinus* in the wild is the result of individual and collective behaviours. While the interflash interval at 1.75Hz is identical for a single firefly or a group of fireflies, the emergence of a well-defined interburst interval necessitates a plurality of individuals. Burst periodicity starts when about 15 males are allowed to interact, a number similar to what was found in previous studies as a threshold for collective behaviour [28]. Each burst consists of a few synchronous flashes, and exhibits a triangular shape (slowly increasing then slowly decreasing number of active fireflies) similar to that observed in the wild, hence confirming that these observations in the wild are true, and not the result of experimental artifacts such as limited depth-of-field. While many fireflies flash in unison during a burst, only a few are flying, while others appear immobile or slowly walking. Finally, male collective display occurs even in the absence of a responding female flash, at least over the course of 15min.

## 5. Method limitations and applications

### (a) Technique validation and limitations

The 3D reconstruction of firefly flashes using pairs of 360-cameras, reported here for the first time (to the best of our knowledge), appears to be generally very reliable and accurate. Reconstructed swarms in the wild follow precisely the geometry of the surrounding terrain (figure 2). Recordings in controlled experiments also faithfully reproduce the shape of the confining tent (figure 5). Spatial streaks and trajectories often exhibit a resolution better than 1cm, as seen in previous figures and accessible to the reader in our interactive FireflyNavigator (ESM 1.2). As with any experimental technique, however, this method has limitations which ought to be acknowledged. We discuss briefly the most significant ones here.

#### Artifacts in 360-degree movies

First, recording over an entire sphere necessarily requires stitching different FOVs together. While this is usually performed by commercial software, small stitching discrepancies are known to be largely unavoidable, creating localization “jumps” along FOV edges in rectangular frames. 360-cameras built around more than 2 lenses might provide better stitching. Second, projecting a sphere onto a plane generates stereographic distortions, so that objects near the poles are stretched out and localized with less accuracy. The GoPro Fusion Studio software, however, enables changing the orientation of the projection, so that if important dynamics occur near the pole of a movie, the axes’ origin can be modified to place these events at the equator.

#### Triangulation resolution

The finite resolution on angle estimation in equirectangular frames implies that triangulation becomes less precise with increasing distance and along the cameras’ connecting line. We discuss theoretical limits to resolution in more details in the ESM 6.1. Briefly, assuming a localization precision of 1 pixel in the equirectangular projection, and given that a frame contains over 3000 horizontal pixels in our movies (spanning 2*π*rad), the resolution on azimuthal angles *θ*_1_, *θ*_2_ is 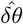 ∼ 10^−3^rad. From geometrical considerations, the error on the distance *r*_1_ to camera 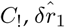, can be related to 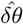 through

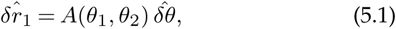

where *A*(*θ*_1_, *θ*_2_) *>* 0 spans several orders of magnitude, and depends on the distance between the two cameras. For our experimental setup, we find that the theoretical triangulation resolution is as low as 1mm in the 1m-radius sphere between the two cameras, and remains below 1cm in a 3m-radius lobe in front of the cameras. Excluding the zones close to the cameras’ connecting line, the theoretical resolution is below 10cm up to 8m away. Increasing the distance between cameras would increase accuracy at large distances, but in a visually occluded environment it would also decrease the likelihood of capturing a same flash in both cameras.

#### Reconstruction results

Regarding our results of stereoscopic reconstructions, we briefly mention the following observations. First, camera pose 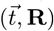 estimations appear very robust across different numerical methods, with discrepancies below 10^−2^ (ESM 6.2). Second, in our swarm reconstructions, about 1% of triangulated points are clear outliers or fall out of physical range, for example far outside the tent volume in our controlled experiments (ESM 6.2). This may be due to improper pairing, or rare occurrences in which two flashes from different sources appear at locations that are compatible in terms of triangulation. Similar problems occur with regular stereoscopic vision, and are better addressed through post-processing filtering.

### (b) Firefly density estimations

While firefly activity shows variability between successive years due to a variety of factors, most notably temperature and humidity conditions, it is widely suspected that firefly populations are generally declining [29]. Climate change, habitat loss, increasing light pollution, and degrading environment are some of the most probable causes. Therefore, estimating firefly densities is fundamental to understanding firefly resilience and promoting conservation efforts [29]. The use of stereoscopic 360-camera setups to record flashing displays is accurate, simple, and inexpensive, and therefore may be appropriate for large-scale monitoring programs. Here, we briefly discuss how 3D-reconstructed data could be used to estimate firefly density. The goal is to estimate the number *N*(*d*) of flashes recorded within a certain distance *d* from the midpoint between the two cameras 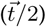. If fireflies were homogeneously spread out in space, *N* (*d*) would grow as *d*^3^. However, for *P. carolinus* at least, we have shown in figure 2*f* that fireflies stay mostly near the ground (a surface), so that *N* (*d*) should actually be expected to increase as *d*^2^. That would be accurate under the ideal conditions of a perfectly-sensitive camera and a bare environment, but in reality the cameras’ limited light sensitivity and visual occlusion from vegetation significantly reduce the number of flashes that can be detected at large distances. Consequently, we expect *N* (*d*) to grow as *d*^*γ*^, with the scaling exponent *γ <* 2. In figure 10, we present the cumulative distribution *N* (*d*)*/N*_total_ as a function of *d* in a log-log plot, and focus on the local slope which indicates the value of the scaling exponent. We find that for *d <* 1m, *γ* ≃ 3, which is consistent with the fact that the considered volume lies within the 2m-layer above ground in which fireflies swarm. For *d >* 10m, *γ* ≃ 0 as such distances are beyond the cameras’ light limitations. But for *d* between 1m and 10m, the scaling exponent is smaller than 2, and closer to 1, indeed reflecting the effect of visual occlusion as discussed above. While it is beyond the scope of the article to propose a complete framework to estimate firefly densities from 3D-reconstructed swarms, the plot in figure 10 could serve to establish a calibration curve to extrapolate large-scale densities from local and imperfect measurements.

**Figure 10.**
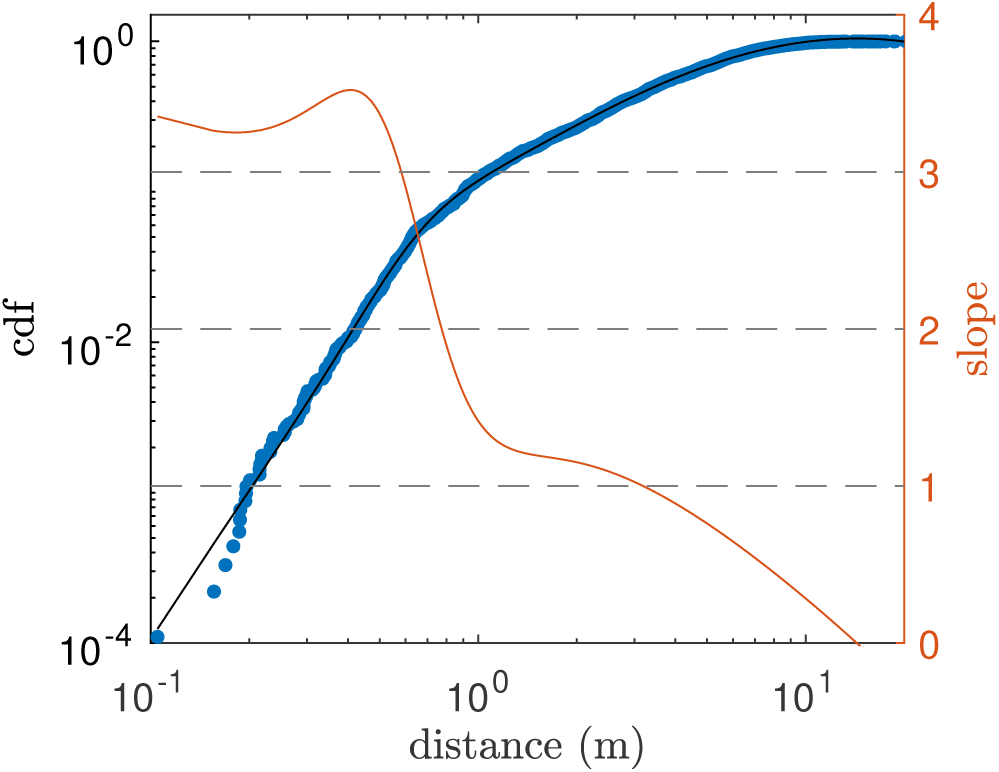
Cumulative distribution function (cdf) of the number of recorded flashes within a distance *d* from the cameras’ midpoint, in log-log plot. In order to approximate local values of the slope, *i*.*e*., the scaling exponent *γ*, a fit of the empirical curve (rational function) was performed (black line), and its derivative is plotted in red on the right axis.

## 6. Discussion

By using pairs of 360-cameras in a stereoscopic setup, we were able to capture the flashing display of *P. carolinus* from within the swarm in a densely forested and visually occluded environment. Triangulation of flash occurrences permitted the 3D reconstruction of mating swarms with sub-centimetric precision within a few meters from the cameras, and hence the identification of specific trajectories consisting of several flashes. A systematic classification of firefly trajectories using statistical methods could provide a basis for the quantitative modelling of behaviour [30]. Our recordings in the fireflies’ natural habitat were complemented by controlled experiments in which a small number of *P. carolinus* males were introduced in a large tent (large enough to allow flying) in order to study interactions between a small number of flashers. Our results in the wild extended prior studies of the intermittent synchrony of *P. carolinus*, and provided additional results relative to the kinematic of firefly trajectories, showing notably different stereotypical flight patterns. Spatiotemporal correlations reflect the flash-burst mechanism of display, and further indicate that instantaneous correlations span several meters, suggesting long-range interactions or a mixture of length-scales. Our most surprising findings come from controlled experiments. We showed that while a single, isolated firefly flashes with a regular interflash interval, its bursts have no periodicity. Only when several fireflies are allowed to interact does a well-defined interburst frequency appear, which suggests that intermittent synchrony is an emergent property of collective behaviour. Controlled experiments also tend to show a differentiation between early and mobile flashers, and immobile followers.

These experimental results will inform future mathematical models that account for species-specific discontinuous flash patterns, long-range spatial correlations, and spatial mixing due to movement of individuals within the swarm. In the meantime, the low cost and implementation simplicity of the 3D reconstruction technique presented here could foster its deployment for large-scale studies of firefly patterns and monitoring programs of firefly populations.

## Data accessibility

The calibration, matching, and triangulation code for pairs of 360-cameras is made available at http://www.github.com/rapsar/stereo360. We are providing two datasets of reconstructed swarms, one in the wild (5min recorded at 30fps on June 5^th^, 2019), and one in the tent with 40 fireflies (60fps). They can be inputted into the FireflyNavigator tool available at https://www.github.com/elie-s/FireflyNavigator.

## Authors’ contributions

R.S., J.H., and O.P. designed and performed the research; R.S. analyzed the data; E.S. designed the FireflyNavigator software; R.S. and O.P. wrote the paper.

## Competing interests

The authors declare that they have no competing financial interests.

## Funding

This work was supported by the BioFrontiers Institute at the University of Colorado Boulder.

## Acknowledgments

We are very grateful to Great Smoky Mountains National Park and the National Park Service for granting us a research permit to observe and record the flashing display of *P. carolinus*. We would like to thank in particular Becky Nichols, the park entomologist, for her guidance, as well as science coordinator Paul Super. In preparing for this experimental campaign we also greatly benefited from Lynn Faust’s previous observations and studies on *P. carolinus*, and her answers to our many questions. Finally, we acknowledge Christoffer Heckman’s helpful advice regarding 3D reconstruction methods, members of the Peleg lab, and Chantal Nguyen in particular, for insightful discussions, and the BioFrontiers Institute at CU Boulder for the utmost support.

## Footnotes

Electronic supplementary material is available online

## Notes

https://github.com/elie-s/FireflyNavigator

