## Supplemental Material Text for "Spatiotemporal reconstruction of emergent flash synchronization in firefly swarms via stereoscopic 360-degree cameras"

### Electronic Supplemental Material for: Spatiotemporal reconstruction of emergent flash synchronization in firefly swarms via stereoscopic 360-degree cameras

Raphaël Sarfati, Julie Hayes, Élie Sarfati, Orit Peleg

March 2020

#### 1 Provided Software

##### 1.1 Matlab Code

A set of Matlab functions to perform three-dimensional reconstruction from pairs of 360-cameras is made available at [see data availability statement for temporary link]. These functions mirror those of the Matlab Computer Vision Toolbox for standard stereo-reconstruction. Below is a brief description of our main functions (see code for more details).

- `xy2alpha`: returns the spherical projections $\vec{\alpha}$ of world points given their $(x, y)$ coordinates in equirectangular movie frames (the size of movie frames in pixels needs to be specified);
- `estimate360CameraParameters`: estimates the relative camera pose $(\vec{t}, \mathbf{R})$ given a set of *matched* projections $\vec{\alpha}_1, \vec{\alpha}_2$ (different methods are possible);
- `matchPoints`: finds optimal pairing for two lists of $(x, y)$ points given $(\vec{t}, \mathbf{R})$;
- `triangulate360`: triangulates pairs of matched projections given $(\vec{t}, \mathbf{R})$.

##### 1.2 FireflyNavigator

Our standalone application FireflyNavigator, made in Unity, is available at `http://www.github.com/elie-s/FireflyNavigator`. The repository also contains a short user guide with instructions, and two datasets of 3D reconstructed swarms: one for the swarm in the tent (`xyzt_tent.csv`), and the other for the swarm in its natural habitat (`xyzt_wild.csv`). Length units are meters.

#### 2 Supplemental Movies

- Movie S1: 360-degree movie showing flash occurrences over 1min, corresponding to the recordings of gp1 presented in figure 2. This movie is an enhanced rendering of actual data. While flashes were recorded in absolute darkness, they are shown overlaid onto the corresponding background (recorded at an earlier time) for orientation. Flashes have also been slightly dilated for better visualization, and coloured in yellow. The movie is rendered in equirectangular frames, and can be visualized as a 2D projection or as a 360-degree movie with Windows' *Movies & TV* or in a Virtual Reality Headset.

- Movie S2: movie of reconstructed flash occurrences over 1min, corresponding to the reconstructed swarm presented in figure 2. The movie is played in real time, and the last few seconds all triangulated flash occurrences.
- Movie S3: movie of reconstructed trajectories corresponding to flash bursts in the tent at $n = 15$ presented in figure 9. Coloured dots of triangulated flash positions, and black dots indicate interpolated flight paths. For better visualization, flash bursts are separated by 2s.

### 3 Additional Experimental Results

To substantiate the findings presented in the Main Text, we provide here additional experimental results.

#### 3.1 Firefly Identification

Fireflies introduced in the tent for our controlled experiments were gently captured using insect nets, and placed in small plastic boxes for inspection. We visually verified that they were *Photinus carolinus* males, using GSMNP's specimen collections as a reference (figure S1).

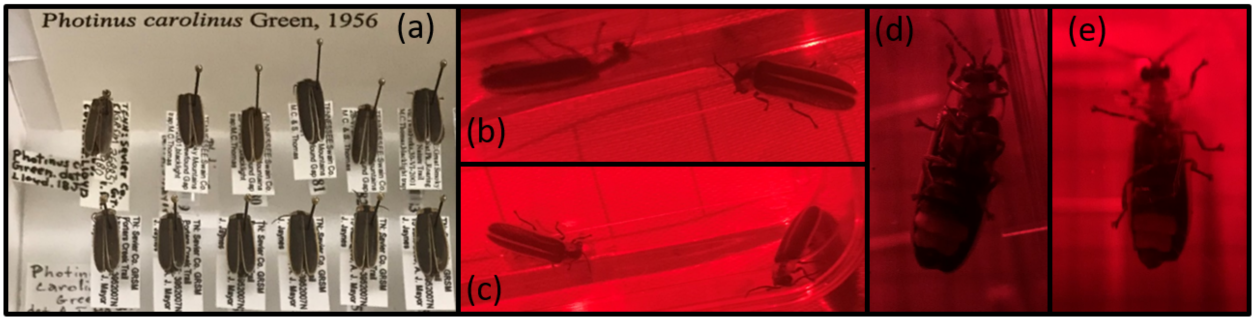


Figure S1: (a) Sample of GSMNP's specimen collection relative to *P. carolinus*. (b,c) Set of pictures of captured *P. carolinus*, dorsal view . (d,e) Set of pictures of captured *P. carolinus*, ventral view.

#### 3.2 Absence of Burst Periodicity at Low Firefly Density

In figure S2 we report additional time series of number of recorded flashes for 1, 2, or 3 fireflies in isolation, from three different sets of controlled experiments. For consistency and clarity, we show intervals of 3min (180s), although experiments typically lasted 5min to 10min. In accordance with the dataset presented in the Main Text, flash bursts do not occur at well-defined intervals, contrary to what is observed when several fireflies are allowed to interact.

#### 3.3 Distribution of Mobilities During Flash Bursts

In the Main Text, we show that during flash bursts in controlled experiments typically only one firefly is flying, while others are standing on the tent's structure (figure 8). Considering the bi-modal distribution of trajectories velocities in controlled experiments (figure S3$a$), it is possible to distinguish slowly-moving trajectories (standing or walking) from fast-moving ones (flying) by setting a threshold at 10cm/s. Based on this distinction, we plot in figure S3 the histograms of the number of flying fireflies per burst in controlled experiments with (b) $n = 15$, (c) $n = 25$, and (d) $n = 40$. Less than half of the bursts have no flying firefly, and among the rest, the majority exhibits only one flyer.

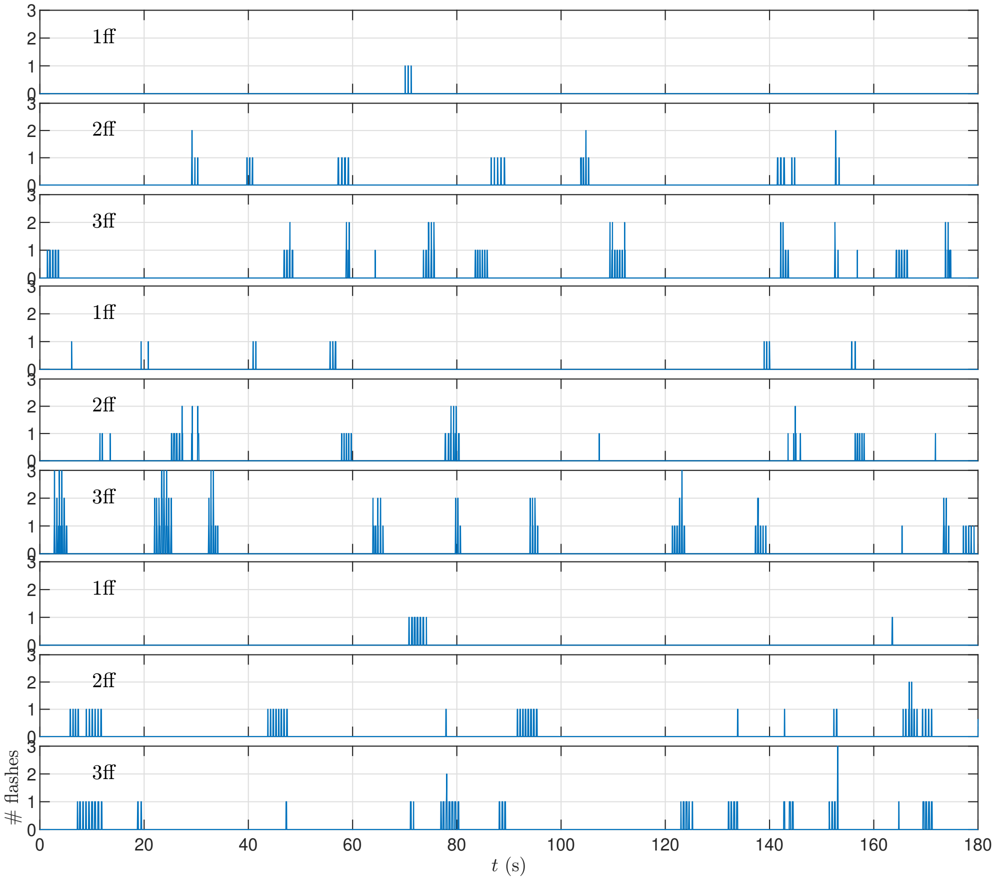


Figure S2: Time series of number of recorded flashes in controlled experiments with 1, 2, or 3 fireflies (ff) in isolation. Despite regular interflash intervals and the onset of synchrony, bursts do not occur at regular intervals of 12.5s as they do at high density of fireflies.

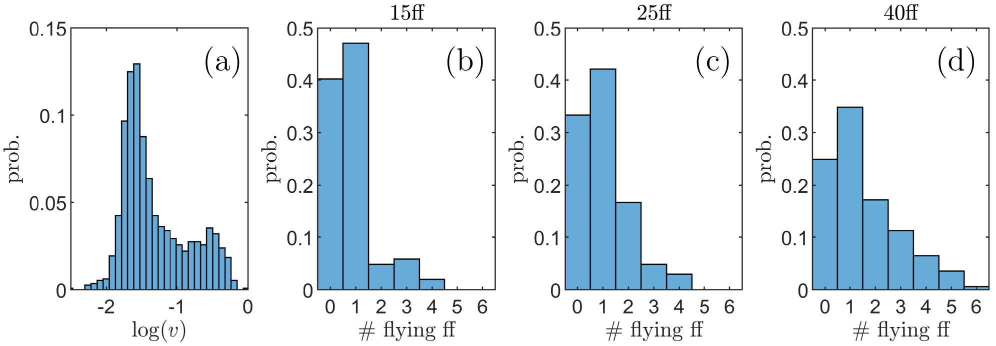


Figure S3: Distribution of mobilities during bursts. *(a)* Distribution of average velocities in individual trajectories. We set the threshold at $10^{-1}$m/s to distinguish between standing and flying trajectories. *(b-d)* Distribution of the number of flying fireflies per burst for *(b)* $n = 15$, *(c)* $n = 25$, *(d)* $n = 40$ff.

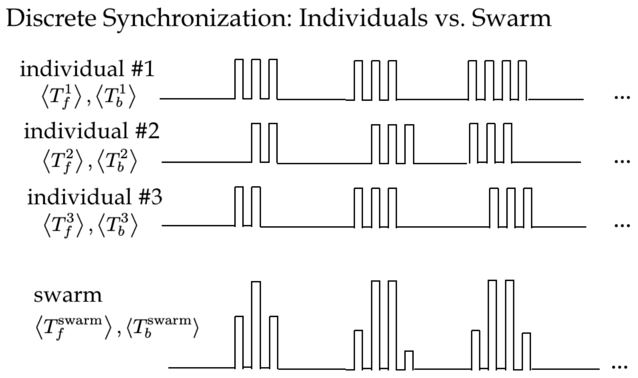


Figure S4: Collective flashing with out-of-phase flash trains leads to triangular flash bursts.

### 4 Temporal pattern of collective flashing

In figure S4, we present a schematic to illustrate the emergence of the triangular shape in bursts of collective flashing. Assuming different fireflies trigger their flash trains at different times, the number of flashes in the swarms over a burst increases then decreases.

### 5 From Positions to Streaks and Trajectories

Individual 3D points can be concatenated into streaks, trajectories, and bursts. Points appearing in consecutive frames are first connected into streaks (spatial localization of a single, continuous flash). This relies on a simple matching algorithm based on minimization of the sum of squared distances within a defined threshold [1].

Based on the flash pattern of *P. carolinus*, it is possible to connect successive streaks into trajectories, notably thanks to their regular interflash interval, between 0.45s and 0.5s [2, 3]. Therefore, for two streaks to be potentially connected, they must occur within that time period. The second condition to connect two streaks is that they are spatially compatible. Since fireflies can be either stationary or fast-moving (see below), this can mean one of two different things. If two streaks occur within a short distance (*e.g.*, 5cm), they are connected. The other possibility to connect two streaks is if they can be linked by a "reasonable" flight path. Specifically, the path between two successive streaks is approximated using a polynomial interpolation (degree = 3) of each spatial coordinate. To be valid, this "likely" flight path must satisfy two criteria: 1) velocities must be consistent with velocities in the recorded streaks; 2) accelerations must be below a certain threshold. Notably, flight paths that would involve high velocities or complicated turns are rejected.

Transitively connected streaks form trajectories. When ambiguities occur, for example a streak potentially connected to two successors, they are resolved by removing edges in the directed graph of streak connections based on a "three-point" algorithm. This means that for two possible 3-node paths going through the same node, the path that will be considered is the one that minimizes the mean acceleration of the interpolated flight.

Finally, a burst is defined as a set of transitively (not mutually) simultaneous trajectories. This is based on the fact that *P. carolinus* collectively flash in bursts separated by a few seconds of total darkness, as detailed below [2, 3]. It is not possible to extend trajectories across bursts, as too much localization information is lost in those few seconds.

### 6 Technique Limitations

#### 6.1 Theoretical Limits to 3D Resolution

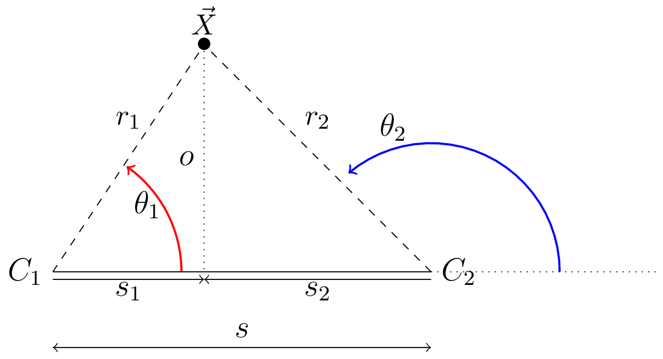


Figure S5: Geometry for triangulation resolution.

To estimate theoretical limits to triangulation resolution, we aim to express the distance $r_1$ between a world point $\vec{X}$ and camera $C_1$: $r_1 = f(\theta_1, \theta_2)$. Then, we assess how small incertainties on $\theta_1, \theta_2$ affect 3D reconstruction. Without loss of generality we consider the upper half-plane, $0 < \theta_1 < \pi$, and triangulation requires $\theta_1 < \theta_2 < \pi$. From the geometrical considerations of figure S4, we have $r_1 = s_1/\cos\theta_1$, $s_1 = 1 + s_2$, $s_2 = o/\tan(\theta_2)$, and $o = s_1 \tan\theta_1$. Consequently, $s_1 = s + s_1 \tan\theta_1/\tan\theta_2$, so that $s_1 = s(\tan\theta_2 - \tan\theta_1)/\tan\theta_2$. Finally:

$$r_1 = \frac{s \cdot \tan\theta_2}{\cos\theta_1 \left(\tan\theta_2 - \tan\theta_1\right)}. \tag{1}$$

Using Wolfram Alpha, we calculate the partial derivatives:

$$\frac{\partial r_1}{\partial \theta_1} = s \cdot \sin\theta_2 \cot(\theta_1 - \theta_2) \csc(\theta_1 - \theta_2) \tag{2}$$

$$\frac{\partial r_1}{\partial \theta_2} = -s \cdot \sin\theta_1 \csc^2(\theta_1 - \theta_2) \tag{3}$$

Assuming now $|d\theta_1| \simeq |d\theta_2| = \hat{d\theta} > 0$, we can approximate the error $\hat{dr_1} > 0$ on $r_1$ as a function of $\theta_1, \theta_2$ as:

$$\hat{dr_1}(\theta_1, \theta_2) = \hat{d\theta} \cdot \sqrt{\left(\frac{\partial r_1}{\partial \theta_1}\right)^2_{\theta_1,\theta_2} + \left(\frac{\partial r_1}{\partial \theta_2}\right)^2_{\theta_1,\theta_2}}. \tag{4}$$

Taking the separation between the two cameras as $s = 1$m and assuming $\hat{d\theta} = 10^{-3}$rad (see Main Text), we solve numerically for $\hat{dr_1}$. The results are presented in figure S6. As expected, resolution is maximum when the rays coming from $C_1, C_2$ intersect at close to 90°; it is minimum when the rays intersect at sharp angles, which is both at large distances, and close to the cameras' connecting line.

#### 6.2 Experimental Reconstruction

**Calibration** Starting from a set of matched points, we propose and implement two general methods for estimating the camera pose $(\vec{t}, \mathbf{R})$: optimization search, and fundamental matrix computation (see Main Text). Fundamental Matrix computation uses a Matlab built-in function, which can

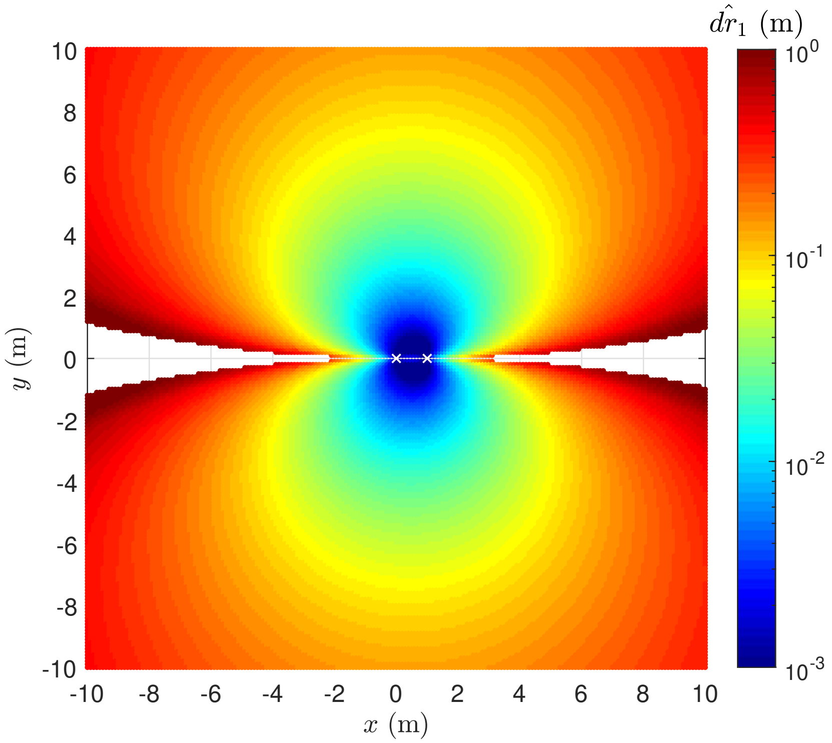


Figure S6: Theoretical limits to 3D reconstruction accuracy, assuming $s = 1$m and $\hat{d\theta} = 10^{-3}$rad. Colours indicate the resolution $\hat{dr}_1$ on the triangulated distance $r_1$, ranging from 1mm (dark blue), to 1m (dark red). The cameras are assumed positioned at $(0, 0)$ and $(1, 0)$ (white crosses).

rely on different algorithms, such as Random Sample Consensus (RANSAC), Normalized 8-Point (Norm8), etc. We show here that the choice of a particular method does not change significantly the results for $\vec{t}$ and $\mathbf{R}$. We use the Fundamental Matrix computation using RANSAC as a reference and compare the discrepancies with the results from other methods in terms of $|\vec{t} - \vec{t}_{RANSAC}|$ and $|\mathbf{R} - \mathbf{R}_{RANSAC}|$ (matrix 2-norm), using the same calibration set of only 17 pairs of matched points (calibration trajectory from a small LED). Further, we show that we can obtain good calibration results by using instead a set of frame pairs in which only a single flash is recorded. We present the results in figure S7. Maximum discrepancies are of the order of $10^{-2}$.

**Triangulation Inaccuracies** When performing the 3D reconstruction of recorded flashes, we noticed some obvious inaccuracies, despite a generally very reliable technique. We briefly mention them here. We also insist that similar inaccuracies can occur in traditional stereo-reconstruction.

First, in the reconstruction of flashes in the tent, we observed about 1% of points outside the volume of the tent (figure S8*a*). These points, curiously, are aligned on two specific planes connecting the cameras to the edges of the tent's curved roof. We believe these anomalous triangulations occur for flashes that take place in those narrow corners, and which may be reflected on the tent's fabric, hence changing their detected localization from their real localization.

Second, in field experiments, we observed that streaks reconstructed far from the cameras tend to exhibit a "seeping" effect, for which streaks tend to align with the direction leading to the camera (figure S8*b*). This is a direct consequence of deteriorating resolution at large distances, as described above.

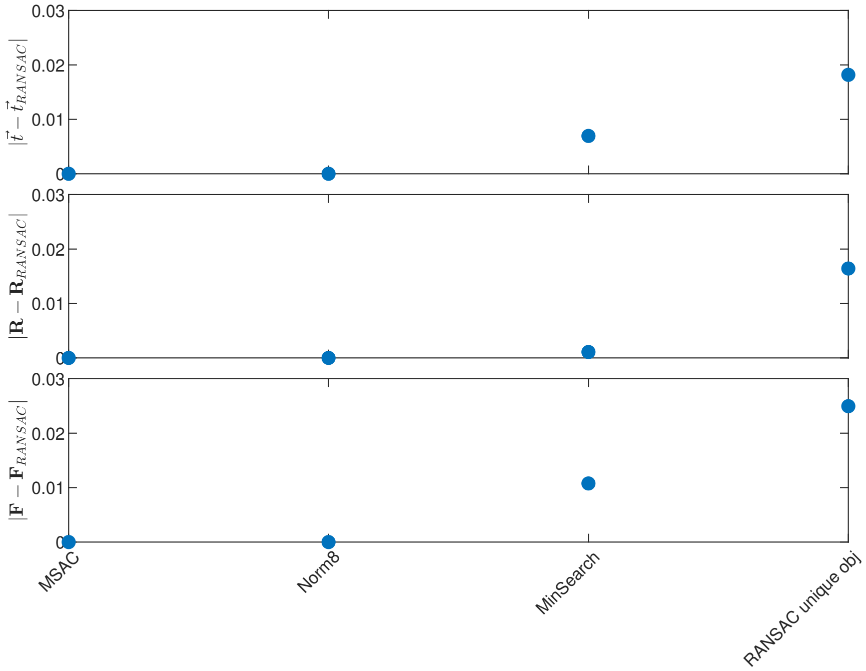


Figure S7: Robustness of calibration against various methods. Estimation of the camera pose ($\vec{t}_{RANSAC}, \mathbf{R}_{RANSAC}$) from Fundamental Matrix computation using RANSAC based on 17 pairs of points from a calibration trajectory is taken as reference. Results from a M-estimator SAmple Consensus (MSAC) or Normalized eight-point (Norm8) algorithm are compared against this reference, along with the optimization search method (MinSearch). Finally, we also compare the results from RANSAC applied to pairs of points coming from single flash occurrences (RANSAC unique obj).

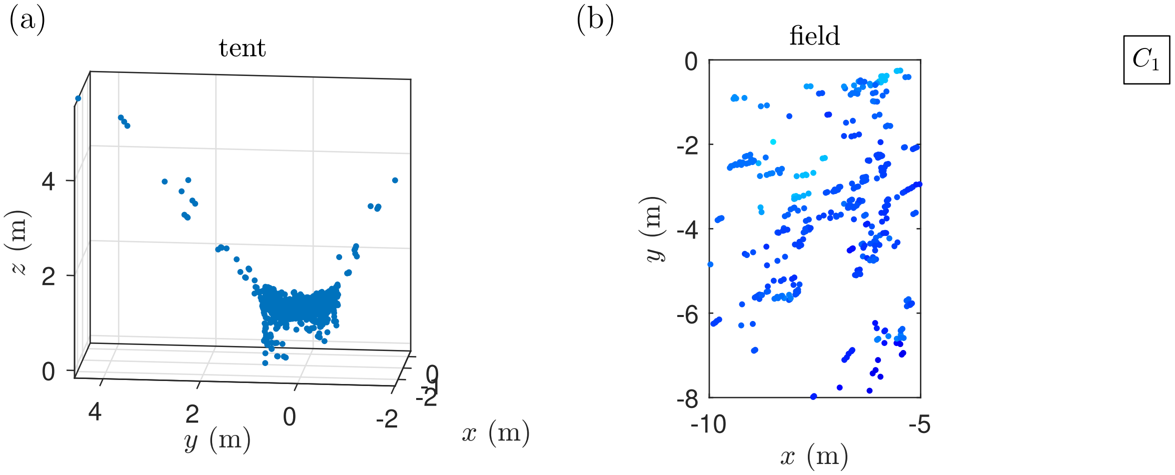


Figure S8: *(a)* Reconstruction of all flash occurrences in the tent experiment. About 1% of triangulated points lie far outside the tent's volume, and along two particular directions corresponding to the curved roof's edges. *(b)* Close-up of the reconstructed swarm from Main Text figure 2$e$, and relative position of one camera ($C_1$). Due to limited resolution, the streaks appear aligned with the camera's direction.
